# Female choice can explain the fertilization pattern in oaks

**DOI:** 10.1101/2021.10.25.465709

**Authors:** Katharina Schneider, Kateřina Staňková, Joel S. Brown

## Abstract

We extend a two-step lottery model of Craft et al. to test the hypothesis that oak trees pursue a form of within-flower female choice to increase the diversity of fathers. Oak trees produce six ovules per flower while maturing just one acorn. When assuming a random ovule selection - which is a natural assumption in the absence of other hypotheses - observed fertilization patterns in oaks cannot be explained: long-distance fertilization is unusually common, even as nearby oak trees may be absent as pollen donors. Our model demonstrates how producing multiple ovules per flower permits selection for rare, distant fathers. The number of ovules per flower that maximizes paternal diversity increases with the number of trees. We introduce a cost function for ovule production for which six ovules per flower balance these costs with the benefits of diversifying fathers. Using data from two published field studies, 7 of 8 investigated maternal oaks had actual paternal diversity indices (average diversity index of 15.42) that fit the female choice hypothesis (estimated diversity of 14.66) significantly better than assuming a random selection from the six available ovules (estimated diversity of 7.649). A third field study permitted us to compare paternity by distance classes for two maternal trees. Both fit the female choice model better than random ovule selection.

## Introduction

In oak trees, distance fertilization - even from outside of the stand - is very common. Using microsatellite markers to study paternity in oaks, Dow and Ashley found more distant pollen donors than would be expected from pollen flow [1,2]. Many acorns were fathered by oaks far from the maternal tree, even as nearby trees fathered remarkably few acorns [2]. However, as in other wind-pollinated species, the likelihood that pollen arrives at a maternal tree decreases with distance to the pollen donor. Thus, there needs to be a mechanism that favors distant pollen donors, because otherwise, the long-distance fertilization cannot be explained. The question “Why are oak trees fertilized by pollen from hundreds meters away rather than from pollen of their direct neighbors” could not be answered convincingly so far [3].

Oak trees are well-known to produce more ovules per flower than will mature into seeds [4,5]. Thus, it seems logical to hypothesize that the overproduction of ovules may somehow lead to the observed pollination patterns. Most species of oaks in the genus *Quercus* have exactly six ovules per flower [4–6], are wind-pollinated, and self-incompatible. Most if not all of a flower’s ovules get fertilized [7], but see [8] and [5] for contradicting studies. In oaks, just one of these six ovules matures into an acorn. Other species that produce surplus ovules include various tropical trees [9–11], *Prunus dulcis* (almond tree) [12], *Phaseolus coccineus* (bean) [13], and *Cryptantha ava* [14].

The overproduction of ovules may be in response to pollen limitation [15, 16], or it may insure some target number of seeds per fruit or pod. Some fertilized ovules may fail as a consequence of genetic defects or post-fertilization genetic load [16, 17]. Alternatively, by way of bet-hedging, the plant may abort some fraction of each flower’s fertilized ovules based on variation in resource availability. However, then, a plant can achieve the same result by aborting flowers rather than ovules within flowers, and such fruit abortion occurs commonly among plant species.

However, these hypotheses may only partly explain observed fertilization patterns [17]. It is unlikely that differences in pollen availability explain why self-incompatible perennials have lower ovule fertilization rates than self-incompatible annuals [18]. Other studies have found that seed-set does not necessarily increase with pollen augmentation [19,20].

Craft et al. proposed that pollinating several ovules while maturing just one permits a form of female choice that increases the diversity of fathers among the acorns of a given oak tree [21]. We assume trees benefit from diversifying the number of fathers among their acorns. Due to a higher diversity of fathers, some are overall more fit than others [22,23], or the increased genetic variability of acorns may be a hedge against environmental contingencies [24–27]. A diversity of fathers reduces sib-sib competition by increasing the likelihood that acorns are half-rather than full-sibs [28, 29]. Finally, fertilization by trees from far away may reduce inbreeding [30, 31].

Craft et al. hypothesized a two-step lottery taking place in the fertilization process of oaks: In the first step, each of the six ovules becomes fertilized [21]. The likelihood of being the father of an ovule is simply a random selection from the pollen grains arriving at the flower. A father providing twice as much pollen to the flower as another has twice the likelihood of pollinating the ovule. Each ovule becomes pollinated in this manner, independent of the other ovules in the flower. In the second step of the lottery, the flower selects the ovule to become the acorn that has no paternal sibs within the flower. If it is not a unique ovule, a random draw from the ovules without paternal sibs selects the surviving ovule. If all ovules have paternal sibs, the surviving ovule is randomly drawn from the ovules with the least number of paternal sibs within the flower. In the remainder of this article, this second step of the lottery is referred to as the female choice. This two-step weighted lottery system increases the odds that more distant oak trees sire acorns. While in the first step, nearby oak trees are more likely to fertilize ovules, their pollen becomes selected against during the second-step due to the presence of more ovules fertilized by these nearby trees. While a simple model of randomly selecting one of the fertilized ovules could not explain the distribution of an oak tree’s fathers, the female choice hypothesis can explain empirical data.

Here, we extend the spatially implicit model of Craft et al. in two novel directions. First, we consider any number of oak trees with a spatially explicit distribution. This becomes essential for comparing model output to actual data. Second, Craft et al. only considered a single focal tree as the mother. Oak trees are non-selfing hermaphrodites [1,2,32,33]. Thus, in our model, we let all trees contribute both pollen and ovules. Our model tracks the diversity of fathers for each oak tree within a stand, as well as the diversity of maternal oak trees among the acorns fathered by each oak tree’s pollen. We validate the model predictions to data from several published data sets that explicitly map the coordinates of trees and their pollen donors.

In the absence of other hypotheses for why oaks produce six ovules and abort five of them, we compare the female choice mechanism to randomly choosing one of the six ovules to mature.

In what follows, we start with a description of our continuous-space model and present simulation results for hypothetical scenarios supporting the understanding of the mechanism. We then compare our model predictions with data from three different published studies [2, 34, 35] to show that the simulations considering female choice in the model explain the data better than a random ovule selection.

## Materials and methods

### 0.1 Continuous-space model with pollen dispersal following an inverse square law

We extend the model by Craft et al. to be able to consider explicit positions of oaks in a continuous space. We apply their two-step lottery model, which determines the father of a flower’s acorn. In step one, the ovules of a flower are randomly fertilized according to each father’s probability *p_i_* to fertilize an ovule of the focal tree. Please note that for simplicity, we assume that all ovules in a flower are fertilized. This assumption allows us to carry out analytical computations. However, in Section 0.3.1, we show that similar results can be obtained when assuming that not all ovules in a flower are fertilized. In step two, based on within flower paternal recognition, each flower “chooses” the ovule from the father that has fertilized the least number of ovules in that flower. If there are six ovules in a flower and three are fertilized by father 1, two by father 2 and one by father 3, only the ovule of father 3 matures into an acorn. All other ovules are aborted. If four ovules are fertilized by father 1, one ovule by father 2 and one by father 3, then either father 2 or 3 will sire the acorn with equal probability. If each father fertilizes the same number of ovules then each has the same probability of siring the acorn. The nearest father has the highest probability of winning step one. However, this father is more likely to lose fertilized ovules during the second step. The opposite holds for more distant fathers as they are favored by the second step.

In contrast to the model by Craft et al., in our model, the probability that tree *i* pollinates an ovule of tree *j* explicitly takes into account the position of *i* and *j* and is given as:

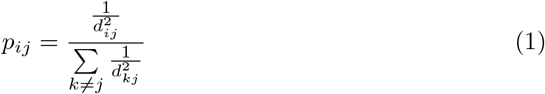

where *d_ij_* is the distance between trees *i* and *j*. Oaks in our model can only fertilize other trees, i.e., we do not consider selfing [1,2,32,33]. All trees function as maternal trees and pollen donors to other trees. We assume that pollen flow follows an inverse square law with distance between any two trees. This assumption approximates the leptokurtic distribution of pollen flow from a tree that has been measured empirically for wind-dispersed pollen [36–38]. For our spatial model, we consider both a torus field with periodic boundary conditions (no edges) and a square field with discrete boundaries. In the remainder of the manuscript, we will refer to these two fields as torus and square, respectively. On a torus, pollen leaving the field on the left/bottom side enter the field again on the right/top side. No tree experiences any boundary effects. The square field scenario imagines woodlots with distinct boundaries. Trees near the boundaries will, on average, be farther away from other trees than trees near the middle.

We use a version of Simpson’s diversity index (SDI) to measure the consequence of the two step lottery for diversifying fathers among an oak’s acorns [39]:

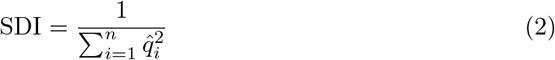

where 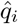 is the fraction of flowers fertilized by father *i* with 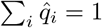 and *n* the number of oaks in the stand. Parameter 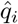 is an approximation for the probability *q_i_* of father *i* winning the two-step lottery in a flower. The SDI lies in the interval [1, *n*]. If each father fertilizes the same number of acorns then SDI = *n*, the total number of fathers. SDI = 1 if only a single father fertilizes all of the acorns of the focal tree.

### 0.2 Model Analysis

In this section, we analyze the relationship between the probability *p_i_* for father *i* fertilizing an ovule of a certain flower and the probability *q_i_* to be the father selected in that flower. Therefore, we here focus on just one single flower. The ovule selection in the flower is based on the female choice mechanism.

The within-flower female choice can skew acorn paternity patterns away from what would happen if there was random ovule selection within a flower. Under random ovule selection, the probabilities *q_i_* to be father of the acorn will conform to the probabilities *p_i_* to fertilize an ovule in the flower, because the expected number of ovules fertilized by father *i* is *ap_i_* with *a* being the total numbers of ovules in the flower and 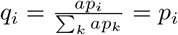. And thus it holds that *p_i_* = *q_i_*. With female choice, the father nearest to the maternal tree still has the highest probability to win the first lottery step. However, this father is more likely to lose fertilized ovules at the second lottery step, and *p_i_* > *q_i_* for trees close to the focal tree. The situation is reversed for the farthest father. He fertilizes the fewest ovules in the first lottery and sees the fewest discarded ovules in the second: *p_i_* < *q_i_* for distant trees. The second lottery favors more distant fathers at the expense of closer ones.

We can derive explicit expressions for the two-stage lottery when there are two or three potential fathers for a focal oak’s flowers. Let *p_i_* be the probability of father *i* fertilizing this particular ovule. We assume that ovules are only fertilized by other trees within the stand and that all trees in the stand contribute pollen that may fertilize ovules of other trees. Thus, 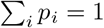. Furthermore, let *X_i_* be the number of ovules fertilized in the flower by father *i*. We then can calculate the probability *q*_1_ of father 1 winning the second lottery (so fertilizing the flower) when considering two fathers in total and six ovules (see Appendix A). This becomes a combinatorial problem. Father 1 only wins the second lottery step for sure if he fertilizes all ovules in that flower, or at least one ovule and at the same time less ovules than father 2. Considering six ovules per flower, this is the case if father 1 fertilizes either one, two, or six ovules. Furthermore, father 1 has a 50% chance to become the father of the acorn, i.e., to win the second lottery step, if he fertilizes the same number of ovules in that flower as father 2. Then, both fathers fertilize three ovules in that flower and the ovule to become an acorn is selected randomly.

Fig 1 shows the relationship between *p*_1_ and *q*_1_ considering two possible fathers and six ovules per flower. In the area above the red dotted horizontal line, it holds that *q*_1_, the probability for father 1 to win the second lottery step, is higher than 0.5. While the red dashed line is showing the relationship between *p*_1_ and *q*_1_ for the random selection mechanism, the blue curve shows the relationship for the female choice model. Considering a random ovule selection (red dashed line), *p*_1_ = *q*_1_. The blue curve, resulting from calculations that can be found in Appendix A, shows the non-linearity in the relationship between *p*_1_ and *q*_1_ for the female choice mechanism. From Fig 1 we see that there are two possibilities for father 1 to have a larger chance than father 2 to win the second lottery (*q*_1_ > 0.5): either *p*_1_ ∈ (0.1, 0.5) or *p*_1_ > 0.9. The first range shows that the rare father indeed has a higher chance of siring a flower’s acorn. However, the rare father needs to fertilize at least one ovule in that flower. Thus, it needs to hold that *p*_1_ > 0.1, because otherwise, father 2 as the common father has a high chance to fertilize all ovules within a flower. For *p*_1_ > 0.9, it is very likely that father 1 fertilizes all ovules within that flower and thus also wins the second lottery step. Fig 1 furthermore shows that the female choice favors rare fathers as the blue curve lies above the red curve for *p*_1_ < 0. 5.

**Fig 1.**
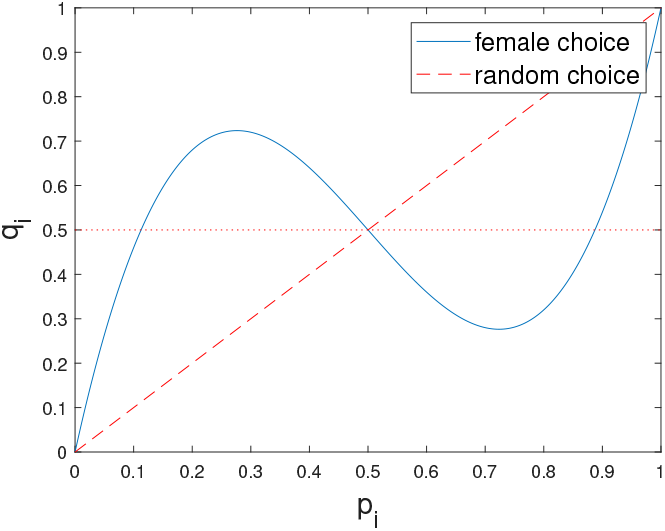
Probability *q*_1_ of a father to fertilize the flower dependent on the probability *p*_1_ to fertilize an ovule in that flower considering *a* = 6 ovules and *n* = 2 possible fathers. The blue curve is derived from Formula 5.

Increasing the number of possible fathers to three fathers makes the calculations more involved. Father 1 will sire the offspring for sure when he fertilizes either all ovules in the flower, or at least fertilizes one ovule but at the same time uniquely the minimal number of ovules in the flower. The following combinations are possible (where the number in the brackets indicate how many ovules are fertilized by the first, second, and third father, respectively): Father 1 either fertilizes six ovules (6,0,0), or one ovule while the other fathers fertilize either none and five ((1,5,0) or (1,0,5)) or two and three ovules ((1,2,3) or (1,3,2)), or two ovules while the others fertilize none and four ovules ((2,0,4) or (2,4,0)). Furthermore, father 1 has a 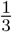 chance to become the father of the acorn, if all three fathers fertilize two ovules ((2,2,2)). If, next to father 1, exactly one other father fertilizes three ovules ((3,3,0) or (3,0,3)), father 1 has a 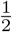 chance to sire the acorn. The calculations can be found in Appendix A.

## Results

### 0.3 Case studies on spatially explicit continuous-space model

We examine the effects of having multiple ovules and the two-stage lottery of female choice within a flower. While having hundreds of ovules is unrealistic for oaks, in our case studies, we consider such high numbers to find optimal numbers of ovules under different environmental assumptions. During the simulations we keep track of the number of acorns each father fertilized. With these values, we are able to compute the SDI according to Equation (2) for each of the trees in the stand. For a given simulation, we then take the average of these SDI values to get a mean measure for the diversity of fathers of a tree’s acorns. Additionally, we can also compute the diversity of maternal trees among the acorns sired by a given tree in the stand. For our simulations, we choose a field of size 10 by 10 units, though the units are arbitrary as Equation (1) is invariant to scale. The random placement of trees creates additional sources of stochasticity compared to the spatially implicit model of Craft et al. Each tree of the stand may have a separate SDI of fathers for its acorns. To account for this, we averaged the SDI across the trees of a simulation. With random placement of trees, the matrix of distances between trees will vary from run to run of the simulation. Thus, we ran 20 simulations for each combination of tree number and ovule number. We give each tree exactly 10,000 flowers.

#### 0.3.1 Simulated SDI for different number of fathers and ovules

Female ovule choice in the two-step lottery can greatly increase a tree’s SDI of fathers compared to a random ovule selection. Fig 2 shows how the average SDI for forest stands with 5, 10, and 20 fathers depends on the number of ovules per flower. The SDI values are averaged over 20 simulation runs and over all trees in the stand and are presented as (*μ* – *s*, *μ* + *s*) where *μ* is the mean and *s* the standard deviation. In all cases, the SDI increases rapidly with the number of ovules up to a global maximum, after which the SDI decreases slowly. Regardless of the number of possible fathers, the maximum SDI is at about 80% of the number of possible fathers. Doubling the number of potential fathers essentially doubles the maximum SDI. However, the maximum SDI occurs at a larger number of ovules per flower as the number of fathers increases. The results are similar whether we assume a torus or a square field. The boundary-less torus reaches a slightly higher maximum SDI, and this maximum is reached with slightly fewer ovules (Fig 2). As would be expected, the variance of the SDI is smaller for the torus than for a square field. Fig 3 shows the results if assuming that only a certain number of ovules per flower (from zero to the number of considered ovules) is fertilized. The number of ovules to be fertilized is hereby drawn from the interval with equal probability. Thus, when considering six ovules per flower, three of them will be fertilized on average. We again observe that the SDI increases rapidly with the number of ovules up to a global maximum after which the SDI decreases slowly. However, the maximum SDI is now slightly higher than when assuming that all ovules in a flower are fertilized. Due to the similarity of these results we assume in the remainder of the manuscript that all six ovules are fertilized.

**Fig 2.**
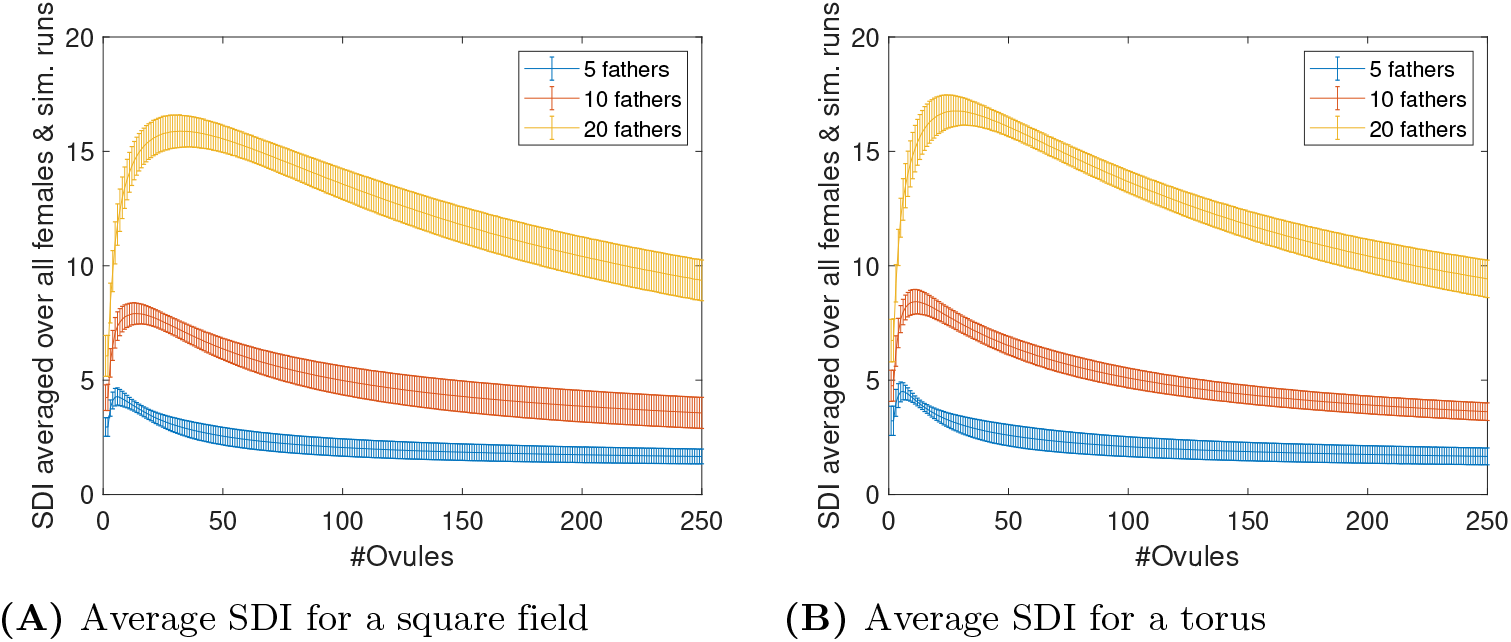
SDI for 5, 10 and 20 fathers on a square (A) and a torus (B) field. Values are averaged over 20 simulation runs and over all trees in the stand. A stand simulated on a square field achieves a lower SDI than the same stand simulated on a torus field. As expected, the SDI on both fields is highest for 20 fathers. However, the SDI follows a similar curve having only one local maximum for 5, 10 and 20 fathers.

**Fig 3.**
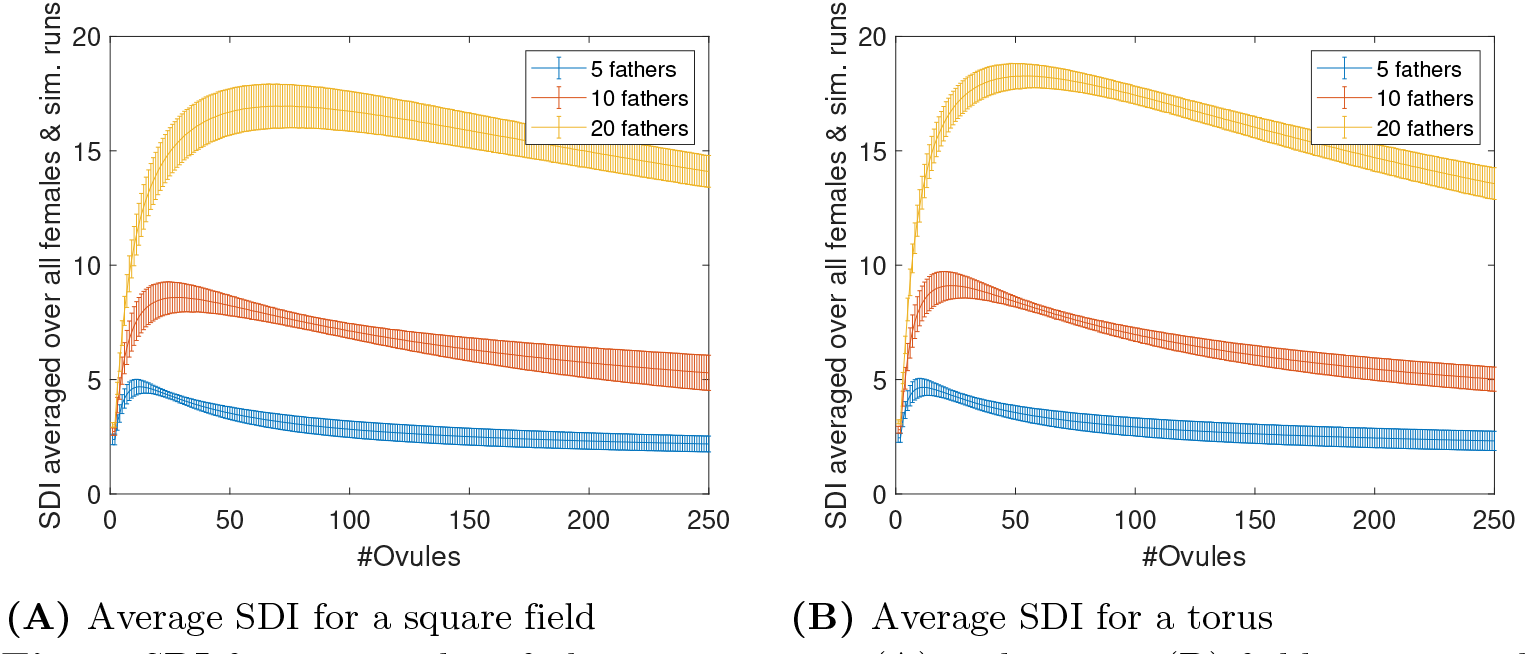
SDI for 5, 10 and 20 fathers on a square (A) and a torus (B) field assuming that not all ovules per flower are fertilized. Values are averaged over 20 simulation runs and over all trees in the stand. A stand simulated on a square field achieves a lower SDI than the same stand simulated on a torus field. As expected, the SDI on both fields is highest for 20 fathers. However, the SDI follows a similar curve having only one local maximum for 5, 10 and 20 fathers.

#### 0.3.2 Optimal number of ovules per flower

For each field (torus and square) and for the different number of trees in the stand (2, …, 41), the values for the SDI maximizing number of ovules per flower are presented as (*μ* – *s*, *μ* + *s*) where *μ* is the mean and *s* the standard deviation of 20 simulation runs (Fig 4). As the number of trees in the stand increases, the optimal number of ovules for maximizing a tree’s SDI of fathers increases almost linearly (Figs. 4A and 4B). This rate of increase is smaller for the stand modeled on a torus than for the stand modeled on a square. For example, to maximize the SDI in an oak stand with 30 possible fathers (31 trees), the flowers would need to have 58 and 43 ovules for the square and torus, respectively. On the square, trees near the border achieve a smaller SDI (see case studies about the preferable position for a tree in a stand). On a torus, no tree experiences such border effects, and thus, the SDI is higher for a torus. Furthermore, the variance in the optimal number of ovules per run of the simulation is higher for the square than for the torus.

**Fig 4.**
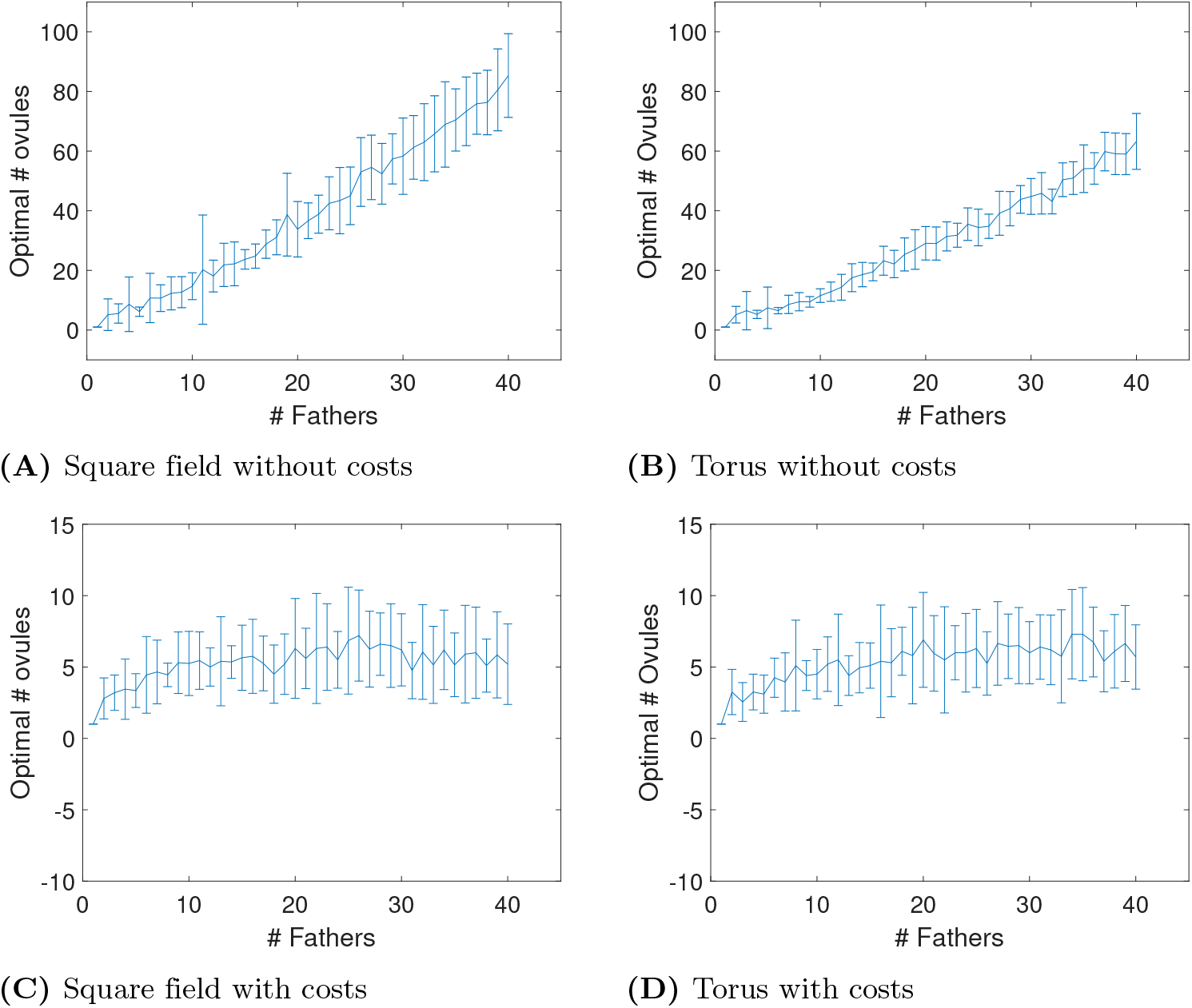
Optimal number of ovules per flower for different numbers of fathers on a square and a torus field. Figs. (A) and (B) do not include costs for ovule production, while Figs.(C) and (D) consider costs for ovule production according to Equation 3. Including these costs, the optimal number of ovules stabilizes around six ovules. For better visualization, (C) and (D) are re-scaled. Without any costs for ovule production, the optimal number of ovules increases linearly with the number of fathers.

However, in nature, we do not observe such large numbers of ovules per flower in oaks but rather six ovules only. It stands to reason that there is a cost to producing multiple ovules per flower. When assuming six ovules per flower in our simulations, we can find costs that lead to six ovules as the optimal number of ovules per flower. For the simulation results displayed in Figs. 4C and 4D, we used costs in units of SDI that increase linearly with the number of ovules produced. As proposed by Craft et al., we reduce the SDI by 0.2 units for each ovule produced, resulting in the following formula for an SDI measure which takes these costs into account, SDI_costs_:

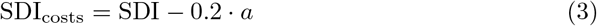

Taking into account these costs for ovule production, the optimal number of ovules increases with the number of trees in the stand and converges to about five to six ovules per flower (see Figs. 4C and 4D).

#### 0.3.3 SDI of stand in relation to the maximal possible SDI

A stand with a large number of trees can achieve a much higher SDI than a stand with a low number of trees. Therefore, to compare the achieved diversity of stands with different numbers of trees, we define the relative SDI_*r*_ by

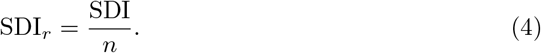

We consider all combinations of numbers of possible fathers in the stand {1, …, 40}, on a square or torus field, and four different numbers of ovules per flower (1, 6, or the optimal number for that number of fathers). We ran 20 simulations of each combination of different numbers of fathers to obtain the mean and variance of SDI_*r*_ (Fig 5). Each simulation re-randomizes the position of trees in the stand.

**Fig 5.**
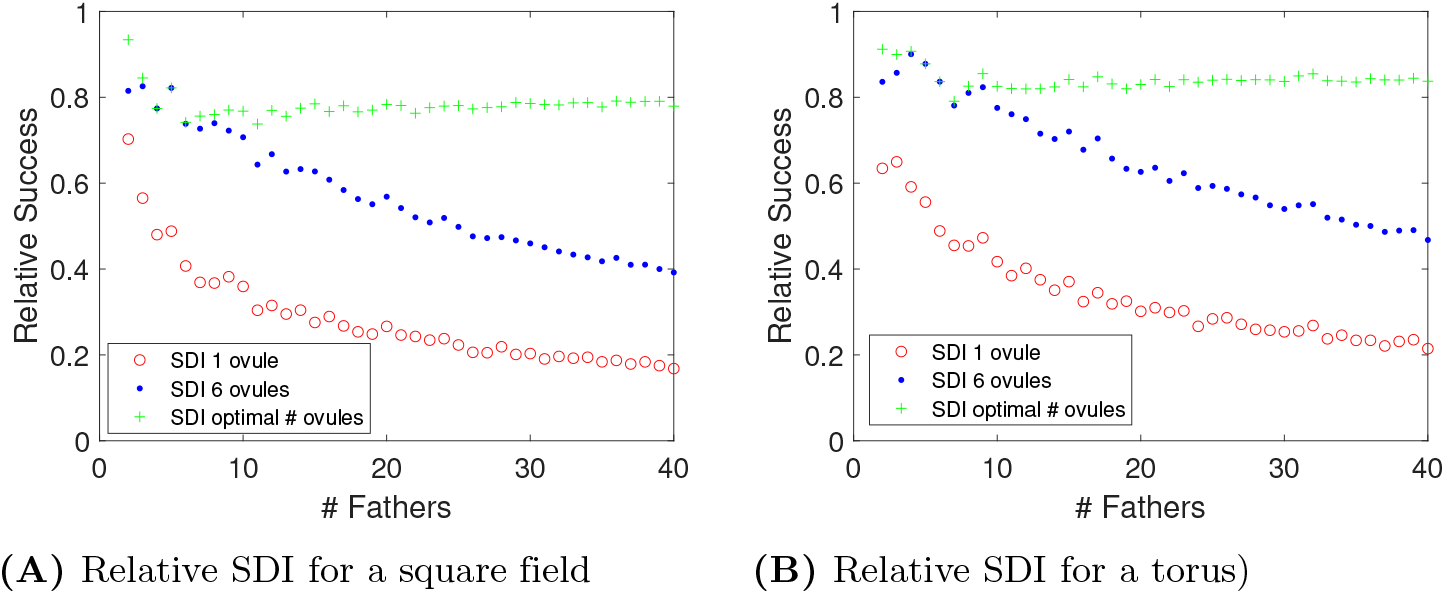
SDI of stand in relation to the maximal possible SDI. The optimal number of ovules is the one maximizing SDI and varies per number of fathers. The optimal number of ovules results in the highest SDI. However, in terms of SDI, having six ovules is a great improvement compared to having just one.

Regardless of the number of ovules, SDI_*r*_ will always start at 100% when there is just one possible father. With just one ovule (no female choice) SDI_*r*_ declines rapidly to around 20% with 40 possible fathers. By definition, SDI_*r*_ is always the highest when the number of ovules is optimal for maximizing the SDI given the number of fathers. It declines from 100% and then flattens out at about 80% once the number of fathers increases beyond four. The empirical value of six ovules per flower is always superior to producing just one. Though, SDI_*r*_ declines steadily with the number of fathers. With 40 fathers, SDI_*r*_ with 6 ovules per flower is between 40 – 50%, substantially higher than for just one ovule. In all cases, the torus generates a slightly higher SDI_*r*_ than the square stand of trees. Given that ovules must come at a cost, we again see six ovules as greatly improving a tree’s diversity of fathers.

#### 0.3.4 Position of a tree in a stand

In a square field, there may be differences between trees near the boundary and those more to the interior of the stand. To examine these boundary effects, we simulated a stand of 1, 000 trees. We evaluated each tree’s acorns SDI of fathers under the assumption of one and six ovules per flower, respectively. We then plotted each tree’s SDI versus its shortest distance to a boundary (Fig 6). There, a vertical and a horizontal line show the mean distance to the border and mean SDI, respectively. With one ovule, the mean SDI (23.56) is less than half of the SDI for six ovules (50.73). At the high end of SDI values, the top 1% of oaks achieved mean SDI values of 113.55 and 130.99 for one and six ovules, respectively. Though, the mean SDI values for the lowest 10% of trees was 2 and 11.5, respectively, for 1 and 6 ovules per flower.

**Fig 6.**
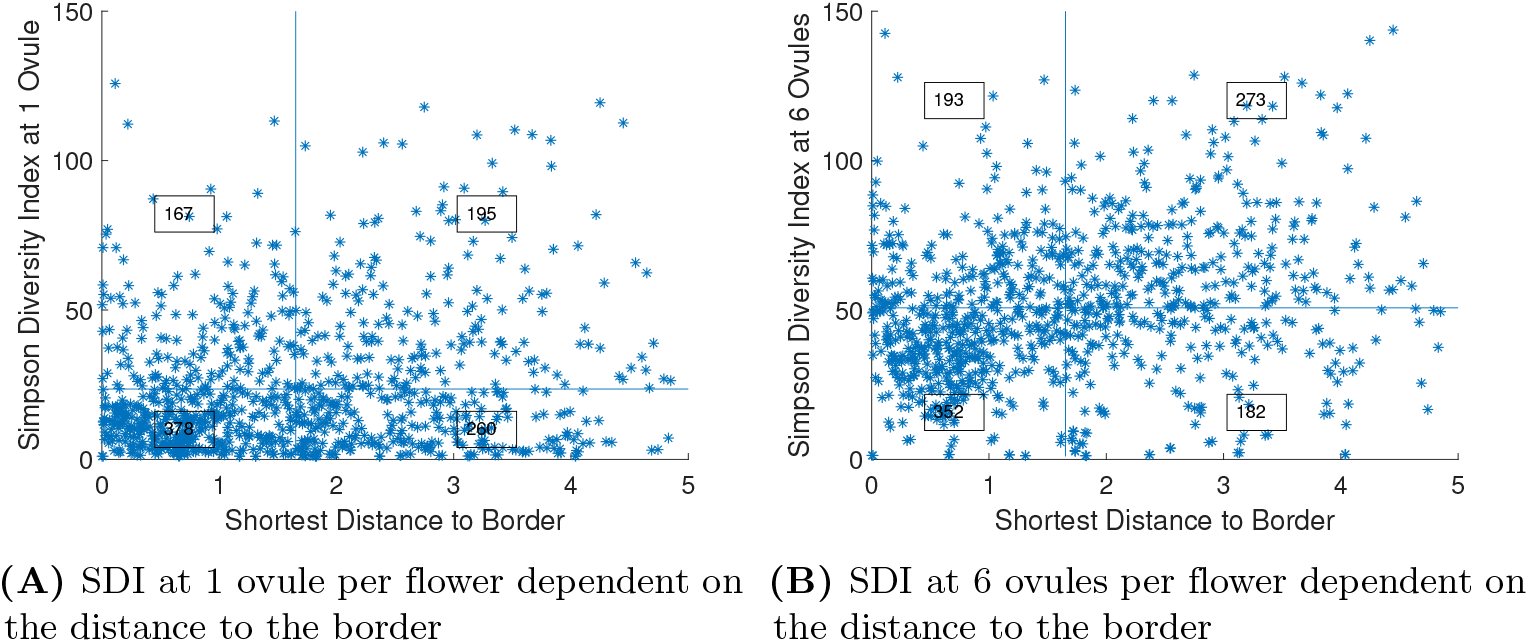
SDI for 1, 000 trees dependent on the distance to the border. The vertical and horizontal lines visualize the mean SDI and the mean distance to the border, respectively. The numbers in the four squares in each figure show how many data points lie within the square. We observe a positive association between distance to the boundary and SDI. With six ovules, the effect is much stronger than with only one ovule.

We used a χ^2^-test on the counts in each of the four quadrants formed by the vertical (distance to boundary) and horizontal (SDI) median lines (Fig 6), to determine whether there is a positive or negative association between distance to the boundary and SDI. Both one ovule (χ^2^ = 16.0, *p* < 0.01) and six ovules (χ^2^ = 16.2, *p* < 0.001) produce a positive association between distance to the boundary and SDI. This suggests that the effective dispersion of nearest neighbors away from the focal tree is higher for those close to the boundary than those on the interior.

### 0.4 Comparing the predictions of one ovule versus six ovule female choice with empirical data

Three published field studies by Dow and Ashley, Pluess et al, and Streiff et al. measured paternity patterns in oak trees in relation to the explicit spatial positions of the trees [2, 34, 35]. We used these studies to compare the predictions between producing just a single ovule versus the six ovule female choice. For generating predictions on the paternity patterns, we used the two-dimensional distance between trees assuming a square field.

#### 0.4.1 Field study of bur oak (Quercus macrocarpa) in Illinois, USA

Using DNA microsatellite markers, Dow and Ashley assigned paternity of acorns collected from three different maternal trees (3E, 17M and 33W) and their pollen donors (see Appendix B, Fig 11) [2]. The majority of the pollen donors were from outside the stand. In these cases, the pollination distance is unknown. Thus, the data used here include the within stand pollination only. From their spatial data, we extracted for each maternal tree how many of the surrounding trees fertilized a particular number of acorns (see Table 1). In this table, the second entry in the second column shows that for maternal tree 3E, 8 trees fertilized exactly one flower. The other entries can be read accordingly. The last rows present the resulting number of acorns where the father could be assigned and the SDI computed per maternal trees (see Equation 2).

**Table 1.** Extracted values from spatial data from a field study conducted by Dow and Ashley [2].

|  | 3E | 17M | 33W |
| --- | --- | --- | --- |
| 1 acorn | 8 | 7 | 10 |
| 2 acorns | 4 | 3 | 1 |
| 3 acorns | 4 | 4 | 3 |
| 4 acorns | 1 | 1 | 2 |
| 5 acorns | 2 | 1 | 0 |
| 6 acorns | 1 | 0 | 0 |
| 7 acorns | 1 | 0 | 0 |
| 8 acorns | 0 | 0 | 0 |
| 9 acorns | 0 | 0 | 1 |
| # acorns | 55 | 34 | 38 |
| SDI | 14.336 | 12.042 | 9.377 |

For each maternal tree, we ran 30, 000 simulations using the spatially explicit distribution of possible fathers. For maternal tree 33W, we made a small adjustment: we did not consider a putative father because its location was nearly identical to maternal tree 33W. As it furthermore did not sire acorns, we concluded that it was probably part of maternal tree 33W. In the simulations, the fathers then fertilized the ovules of a flower where the flower number was set to the actual number of acorns sampled for a given maternal tree. We calculated the tree’s paternal SDI assuming female choice with six ovules (two-stage lottery) versus random selection from the fertilized ovules of a flower. For all three maternal oaks, the SDI for female choice is higher than for random ovule selection (see Fig 7). For tree 3E, the actual SDI lies very close to the mean of the SDI under the random ovule selection, though it lies within the range of possibilities for female choice. For the remaining two maternal oaks (17M and 33W), the actual SDI coincides better with the simulation results for the female choice than for the random selection. For these two trees, the actual SDI is beyond any of the 30, 000 values generated by random selection (Fig 7).

**Fig 7.**
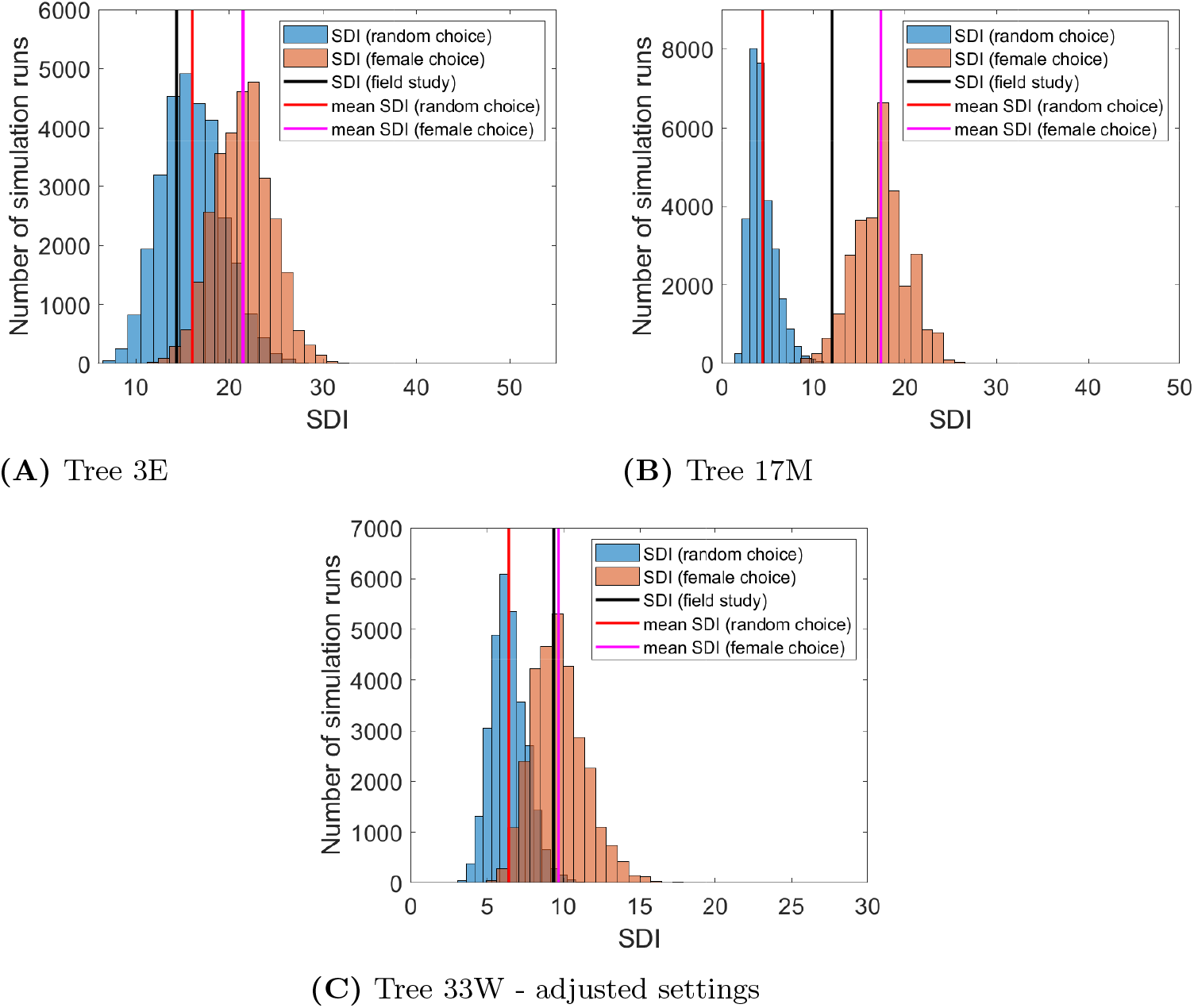
Histograms of the paternal SDI of 30,000 simulation runs. The blue and orange histograms present the values considering one ovule (without female choice) and the values considering six ovules (with female choice) per flower, respectively. For tree 33W (Figure (C)), we dropped a putative father because its location was near identical to maternal tree 33W and hence likely from the same plant. Furthermore, it sired no acorns.

#### 0.4.2 Field study of *Quercus* lobata Nee in the Sedgwick Reserve, USA

Pluess et al. present spatial data on five maternal trees and their pollen donors within a 250m radius (see Appendix B, Fig 12) [35]. In order to compare these results with results obtained by simulations with our continuous-space model, we can compute the SDI for each of the five maternal trees. Pluess et al. do not provide exact numbers of offspring for each of the paternal trees. Rather, they present paternity based on fathers at different distance intervals (see column “Offspring data from field studies” in Table 2). Therefore, we used the middle distance of each interval for our computations (see column “Number offspring used as input for SDI calculations” in Table 2). The number of acorns under investigation varied from 82 to 108 for each maternal tree (see row “Number of acorns”). The resulting SDI values are shown in row “SDI”.

**Table 2.** Extracted values from spatial data from a field study conducted by Pluess et al. [35]

| Offspring data from field studies | Number offspring used as input for SDI calculations | Tree 32 | Tree 35 | Tree 165 | Tree 844 | Tree 889 |
| --- | --- | --- | --- | --- | --- | --- |
| 0-1 | 1 | 22 | 22 | 10 | 19 | 11 |
| 1-2 | 2 | 4 | 0 | 5 | 0 | 2 |
| 2-4 | 3 | 7 | 6 | 11 | 4 | 8 |
| 4-10 | 7 | 3 | 4 | 4 | 3 | 7 |
| >10 | 10 | 1 | 2 | 2 | 4 | 2 |
| Number acorns |  | 82 | 88 | 101 | 92 | 108 |
| SDI |  | 19.3218 | 16.4068 | 19.4305 | 14.0598 | 18.3975 |

Fig 8 shows the histograms for the SDI of 30, 000 simulations when using the exact tree locations of the field study and using the same number of flowers per tree as acorns empirically investigated (see Table 2, row “Number Acorns”). For all cases, the actual SDI conforms more closely to female choice (six ovules) than random ovule selection (one ovule). In two cases, the actual SDI is close to the simulated mean for female choice (one higher and one lower). In the other three cases, the actual SDI values are considerably higher and in the upper tail of the distributions for female choice and far outside any of the simulated values when there is random selection (Fig 8).

**Fig 8.**
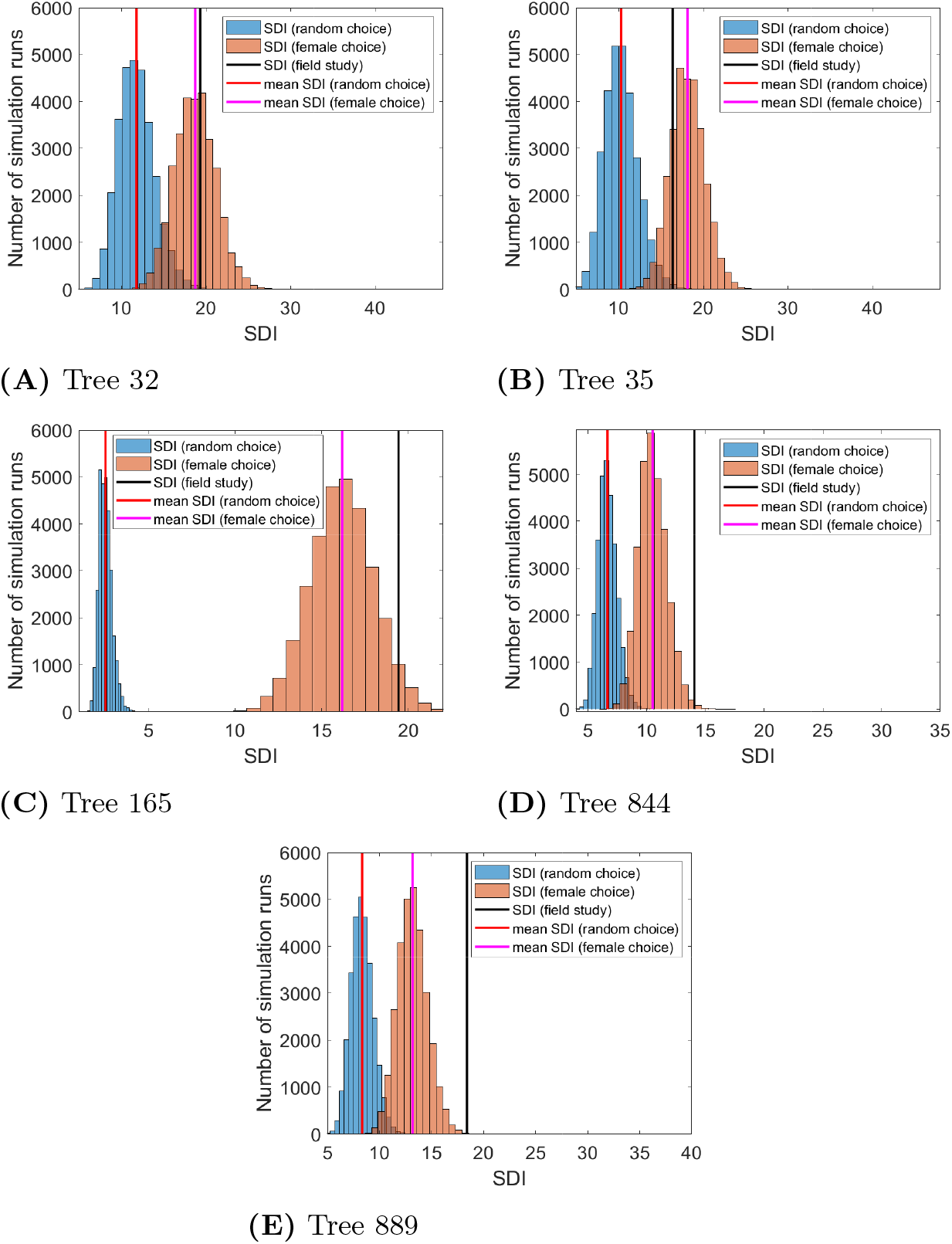
SDI histogram of 30,000 simulation runs for each of the trees 32, 35, 165, 844 and 889 from the field studies by Pluess et al. [35]. The orange histograms contain the SDI values assuming a female choice of ovules and the blue histograms the SDI values considering a random selection from the six ovules of a flower. For all five cases, the simulated SDI values are higher for female choice than random ovule selection.

#### 0.4.3 Field Study of *Quercus* petraea and *Quercus* robur in the northwest of France

Streiff et al. examined the spatial distribution of paternal trees for two focal trees of species *Quercus petraea* (acting as females) in an oak stand in the northwest of France (see Appendix B, Figs. 13A and 13B) [34]. They compared potential and actual paternal trees in relation to their distance to maternal trees within a mixed oak stand of *Quercus robur* and *Quercus patraea*. Potential trees are those trees that could have sired acorns of a focal maternal tree, while actual paternal trees successfully fertilized acorns of the focal tree.

The distribution of potential fathers (blue bars in Fig 9) for maternal tree E increases with distance. This is to be expected for an evenly or randomly distributed stand of trees, since the area covered by a distance ring will increase with distance (see Appendix B, Fig 13). In contrast to that expectation, we see a skewed distribution of trees surrounding maternal tree B. With a maximum at the 50m category, we observe a steady decline in tree numbers at 70m, 90m, and 110m. This happens because tree B exists within a clump of trees that thins with distance. For both trees, nearby trees are the likeliest fathers, though distant trees are well represented among the acorns (red bars in Fig 9).

**Fig 9.**
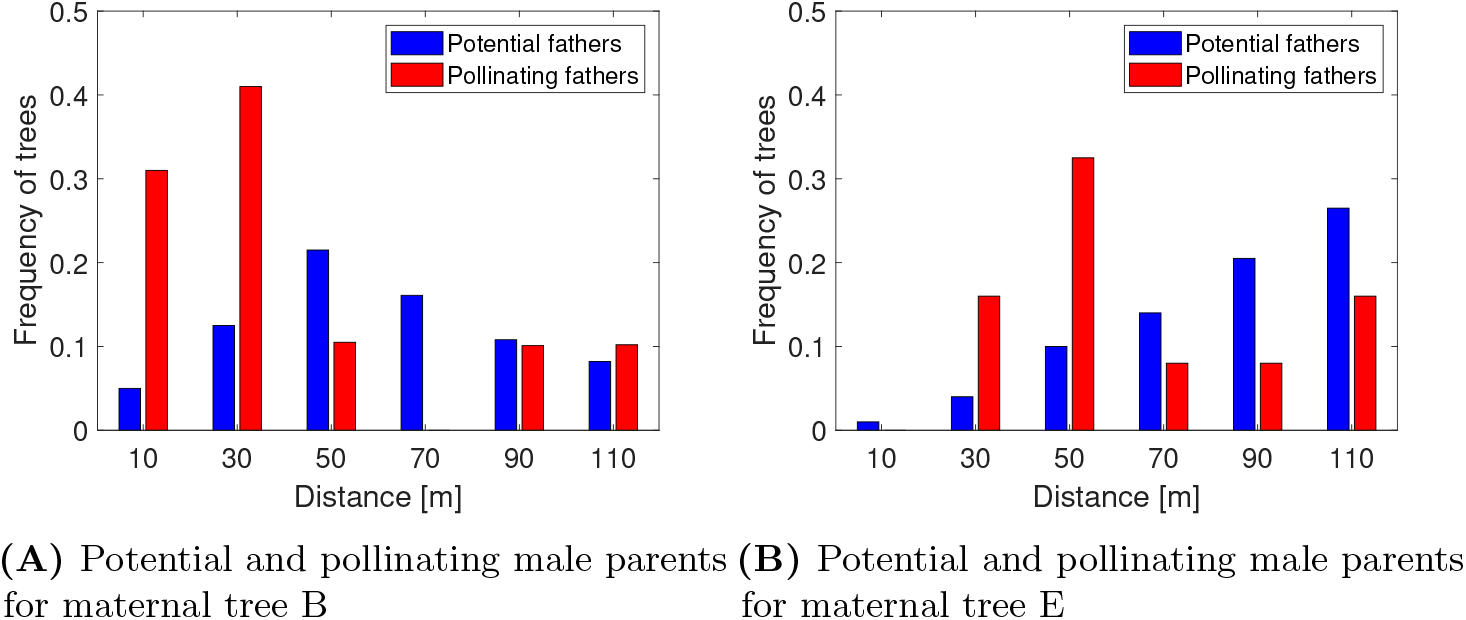
Figs 9A and 9B show a comparison of potential (blue bars) and pollinating (red bars) male parent distributions dependent on the distance to maternal tree B and maternal tree E. The results are taken from Streiff et al. [34].

We performed 30, 000 simulation runs and considered 21 flowers for maternal tree B and 20 flowers for maternal tree E. This coincides with the number of acorns tested for paternity. Because they present paternal information binned into distance categories from the maternal trees, we focus our analysis on the distance-dependent fertilization pattern (Figs. 9A and 9B).

As expected, the female choice model increases the frequency of more distant paternal trees when compared to random selection. In order to evaluate whether the female choice hypothesis better explains the fertilization patterns, we compare the errors associated with these simulations and the actual data. We use the squared differences of the simulated frequencies *x*_sim_ and real data points *x*_real_ as the error measure. For both maternal trees (Figs. 10A and 10B), the average error of 30, 000 simulation runs was lower for the female choice model (Tree B: 0.0257; Tree E: 0.0262) than for the random ovule selection model (Tree B: 0.0449; Tree E: 0.0332).

**Fig 10.**
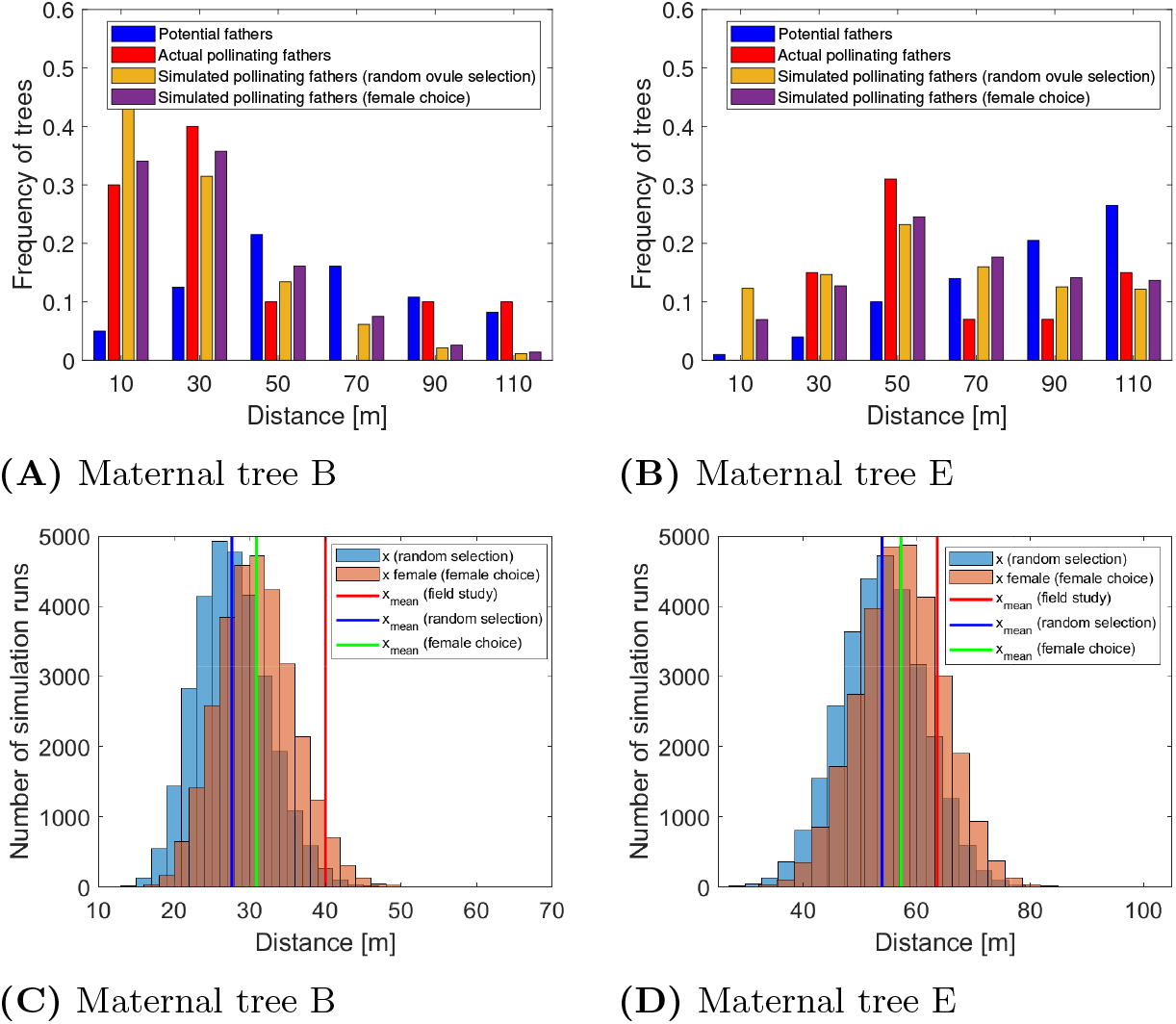
Comparison of data obtained from field studies by Streiff et al. to data obtained from simulations for both maternal trees B and E. Figs. (A) and (B) show the frequency of paternal trees that have fertilized flowers of the maternal tree dependent on their distance classes. In Figs. (A) and (B), the blue bars indicate the potential fathers and the red bars the actual pollinating fathers from the field studies, whereas the yellow and magenta bars show the simulation results considering random ovule selection and female choice, respectively. Figs. (C) and (D) show the distance histograms of fertilizing paternal trees. In Figs (C) and (D), the blue histograms show the mean distances of successful fathers to the maternal tree from 30, 000 simulation runs with random ovule selection. The orange histograms show the distances when considering a female choice. The red line indicates the estimated mean distance of the fertilizing trees from the field study. We approximated it by taking the middle of the distance classes as an estimate for the distance for all trees within this class.

In evaluating the mean distance of fathers from the focal tree, the random selection simulations yielded the smallest mean distance, the female choice model yielded an intermediate distance, and the actual data had the highest mean distance value. For Tree B, both random selection and female choice substantially underestimated the actual mean. For Tree E, the actual value lies close to the mean for the female choice model, and far from the mean for the random selection model. Regardless, the female choice model provides a better fit to the actual data (Figs. 10C and 10D).

## Discussion

Craft et al. hypothesized that oaks produce multiple ovules per flower as a means of the individual tree to diversify the number of fathers across its acorns [21]. We found modeling and empirical support for this hypothesis. For wind-pollinated species like oaks, the challenge is to discourage nearby pollen donors and encourage more distant ones. This can be achieved through a two-step lottery. The second step of the lottery can happen if a plant, such as an oak tree, fertilizes multiple ovules per flower but matures just one seed per flower. Even if ovule fertilization occurs randomly with respect to pollen flow (favors nearby paternal trees), a tree can give an advantage to rare pollen donors by maturing a flower’s acorn from the ovules of a flower that have the fewest fathers. A nearby pollen donor sees its contribution discarded when it fertilizes two or more ovules of a flower. Step one of the lottery strongly favors the nearest paternal trees. Step two favors the more distant pollen donors.

For this flowering strategy to work, the plant must produce three or more ovules per flower. The optimal number of ovules per flower to maximize the diversity of paternal oaks among a tree’s acorns depends strongly on the number and the dispersion of oaks within and between forest patches. Without any costs for ovule production, this optimal number can range from as little as 5 to well over 100; and it tends to increase with the number of potential fathers and with the decline in pollen flow between successively more distant paternal trees. The six ovules characteristic of most oak species achieves large increase in diversity relative to having one. Having six ovules per flower may balance the benefits of diversifying fathers against the costs of producing and discarding multiple ovules within a flower [40]. When assuming six ovules per flower, we found - as a showcase - that there exist costs for which the benefits of diversifying fathers and costs are balanced.

In our case studies, we assumed that all six ovules in a flower are fertilized. However, there are some studies claiming that not all ovules are fertilized [5, 8]. Thus, in Section 0.3.1, we showed that we obtain similar results when assuming that not all ovules in a flower are fertilized.

Ten trees from three separate studies provided information on the paternity of their acorns [2, 34, 35]. From these studies, we evaluated the distribution and diversity of fathers. We compared empirical results to our model of female choice (with six ovules per flower) and to random selection where fathers of acorns occur in proportion to their pollen contribution. In all examined cases, random selection predicted a lower diversity of fathers than within-flower female choice. For nine of ten trees, the data more closely fit the predictions of female choice than random selection. For the nine cases, the female choice model either closely fitted the actual data (6 times) or underestimated the true diversity of fathers (3 times). The model will underestimate the diversity of fathers if there are potential pollen donors beyond the boundaries of the studied stand. The data of the three studies we used to validate our model included only within-stand pollination. Thus, the real SDI in these studies is most likely - as well as our modeled SDI - higher than the SDI that we used to validate our model. In the studies of Dow and Ashley, more than half of the pollen donors come from outside the stand [1]. From the studies of Pluess et al., we used paternal data from at most 250 metres from the maternal tree [35]. In Streiff et al., more than 60% of the pollination was from other than the investigated oaks. In total, this makes the difference of the empirically measured SDI to the modeled SDI obtained under the assumption of a random ovule selection even larger [34].

To our knowledge, no other hypothesis so far can explain the fertilization pattern in oaks. Thus, due to the absence of other hypotheses on how ovules are selected, we here compared the result of applying a female choice mechanism to results when applying a random ovule selection. We showed that the female choice mechanism explains the data better than a random ovule selection. Still, our results do not prove the existence of the female choice mechanism. Our model can, however, be used to validate new upcoming hypotheses.

A number of factors not considered in the present model would likely skew the diversity of pollen donors to a given oak tree. One factor we did consider was how a centrally located oak will have a higher diversity of fathers among its acorns than oaks nearer the boundary of the forest. Other important factors influencing pollen flow and hence paternity (whether random or through the female-choice two-step lottery) include topography and wind properties. Different formulations for pollen flow would expand the utility of the model, though current data may not yet permit model validation for these additional features. Studies showing leptokurtic pollen flows for wind-pollinated plants provide a starting point [37,41]. In future research, it would be interesting to examine how different pollen flow models, such as a multiscale model by DiLeo et al., could further improve the fit of our model [42]. Furthermore, we so far only examined pollination within a closed stand of oak trees. In nature, pollen arrives as well from outside the stand. Taking into account also father trees from outside the stand may change the paternity analysis [43].

Our model considered the trees as points on the landscape. Actual trees occupy space with varied and expansive canopies. Small or less robust trees producing less pollen will be less successful in the first step of the lottery but will have more to gain during the second step. Our female choice model adds another source of diminishing returns to a tree from producing more pollen.

Proximity to different sides of a neighbor’s canopy should matter. One would expect the pollen of a southward neighbor to contribute disproportionately to the south rather than the north side of a focal tree [44]. Data on this is sparse, and data for paternity in oaks has not yet been localized to position on the maternal tree. If nearby pollen donors spatially clump their pollen onto different regions of a recipient tree, then our model of female choice would further enhance the diversity of fathers based on distance. To see this, imagine four trees all close and equally distant to a maternal tree. Assume that they only differ in their orientation to the tree. If these trees contribute pollen randomly across the focal tree, they would mutually favor each other relative to more distant trees. This is because they would reduce the odds of one of their own fertilizing two or more ovules of a flower. Thus, less of their pollen would be discarded during the second step of the lottery. The focal oak would be less effective at discarding the ovules pollinated by nearby trees. On the other hand, if each of these four neighbors concentrates its pollen on the most proximal region of the focal tree, then many more flowers would have multiple ovules fertilized by the same tree. These would be discarded. Aspects of this could be modeled and investigated by giving each tree’s canopy spatial dimensions and letting pollen flow correspond to directionality and proximity.

Other empirical studies have examined the diversity of fathers in self-incompatible plants and how various mechanisms could enhance such diversity. Lankinen and Madjidian found that delayed stigma receptivity in the species *Collinsia heterophylla* results in a higher paternal diversity [45]. Others have modeled and proposed ways for plants to exhibit female choice. Melser and Klinkhamer experimentally tested the hypothesis that females of the species *Cynoglossum officinale* choose to abort low-quality offspring [46]. They investigated the effects of artificially removing ovules and the effects of adding nutrients. They found that offspring survival was lower with random ovule abortion (by hand) than when the plant aborted ovules. As expected, adding nutrients increased offspring survival. Furthermore, Sakai modeled the hypothesis that females choose certain fertilized ovules to create seeds of uniform size [47]. By producing surplus ovules, females can choose ovules that absorb a similar amount of resources.

Our model requires some sort of self/non-self recognition among fertilized ovules of a flower. The flower must recognize full-sibs (same father) relative to half-sibs (different fathers). Studies show that selective abortion takes place in many trees [48,49]. To our knowledge, there have not been studies done looking at the distribution of full- and half-sibs within the fertilized ovules of an oak. We proposed that fertilization is random with respect to pollen accrual to the flower. Yet, other mechanisms exist for preventing successful fertilization of ovules based on mating types or histone compatibility. Self-incompatibility systems genetically prevent fertilization from the plant itself or closely related plants [50–52]. These systems can further be distinguished between gametophytic and sporophytic systems [53]. Moreover, pre-fertilization barriers might prevent fertilization from different species. In several tulip species, Creij et al. identified different forms of barriers that occurred at various developmental phases [54]. Barriers to fertilization could include failure of pollen germination, pollen tube growth, and pollen tube penetration of ovules [54]. These pre- and post-fertilizing mechanisms have been more often examined for short-lived plant species than for long-lived species like oak trees [55]. Hagman et al. showed how species of *Quercus* use a gametophytic control of pollen tube growth to prevent selfing [56]. Furthermore, Boavida et al. investigated post-pollination mechanisms in the species *Quercus suber* [57]. Their study examined pollen-pistil interactions in order to gain insights into intra- and interspecific crosses.

Oaks are not alone in having multiple ovules per flower while maturing just one. In species of *Symphoricarpos* and *Cornus* only one out of multiple ovules mature [58]. Moreover, in *Erodium cieutarium*, only one ovule per schizocarp develops into a seed [58]. We can observe a similar behavior in the species *Pongamia pinnata*, that matures only one of the two seeds in most of its pods [59]. Although, with just two seeds, some sort of between flower selection would be required to improve paternal diversity.

Our model can be extended and made more realistic. Such extensions could be used to predict the distribution and diversity of fathers for a focal plant’s seeds. While the model showed promise, it needs and invites empirical tests of its assumptions, hypotheses and predictions. For instance, natural and controlled pollination experiments with oaks in the field could vary the pollen mix of fathers (either by distance or hand pollination) to subsets of flowers of focal individuals. One could then measure a father’s success from this mix in fertilizing ovules of a flower, and in siring acorns. Such experiments could answer the question of whether fathers contributing less pollen gain a proportional advantage at either the ovule stage or, as predicted, at the acorn stage.

## Acknowledgments

We thank Frank Thuijsman and Ralf Peeters for providing valuable insights, advice and encouragement. We thank Mary Ashley for assisting us in accessing and interpreting the empirical studies. This research was supported in part by a graduate internship at the Moffitt Cancer Center, Tampa, Florida; and by grants from the Transnational University Limburg, and by the European Union’s Horizon 2020 research and innovation program under the Marie Sklodowska-Curie grant agreement No 690817.

## A Appendix A

Let *p_i_* be the probability of father *i* fertilizing a fixed focal tree *j*. We are only interested in which surrounding tree is fertilizing ovules of the focal tree *j* and thus, we assume that only the focal tree *j* is fertilized and that all other trees in the stand do not fertilize each other. Furthermore, let *X_i_* be the number of ovules fertilized in the flower by father *i* and let *a* ≥ 3 be the number of ovules in a flower. We assume that the number of fertilized ovules, 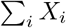, is always equal to the total number *a* of ovules in a flower and *X*_1_ ≠ 0. Furthermore, we assume that 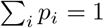.

We then can calculate the probability *q*_1_ of father 1 winning the two step lottery (so really fertilizing the flower) when considering two fertilizing fathers in total and six ovules per flower:

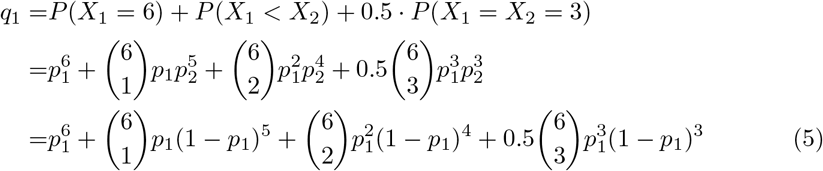

In Fig 1, we display *q*_1_ as a function of *p*_1_ based on this formula.

Let *δ_k_*(*b*) be 1 if *b* is divisible by *k*, and 0 otherwise. When considering 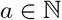 ovules per flower, the probability *q*_1_ for father 1 to fertilize the flower becomes:

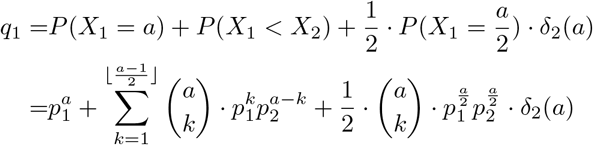

Increasing the number of possible fathers to three fathers (*i* ∈ {1, 2, 3}), makes the calculation for six ovules per flower slightly more complicated:

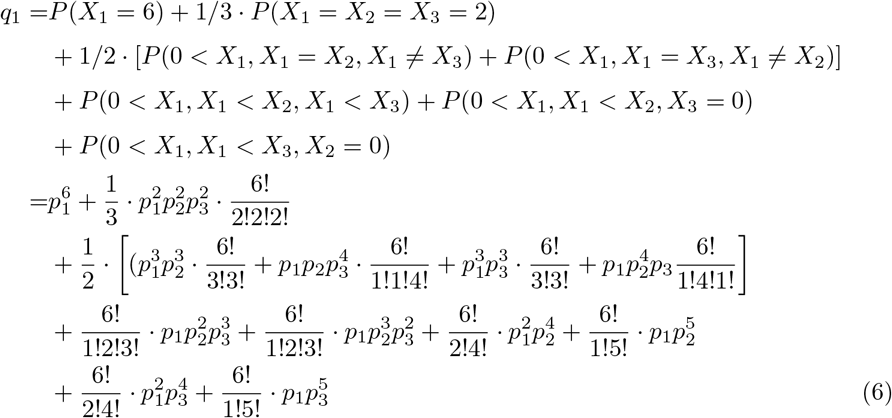

For *a* ovules per flower and still considering three possible fathers, we obtain the following:

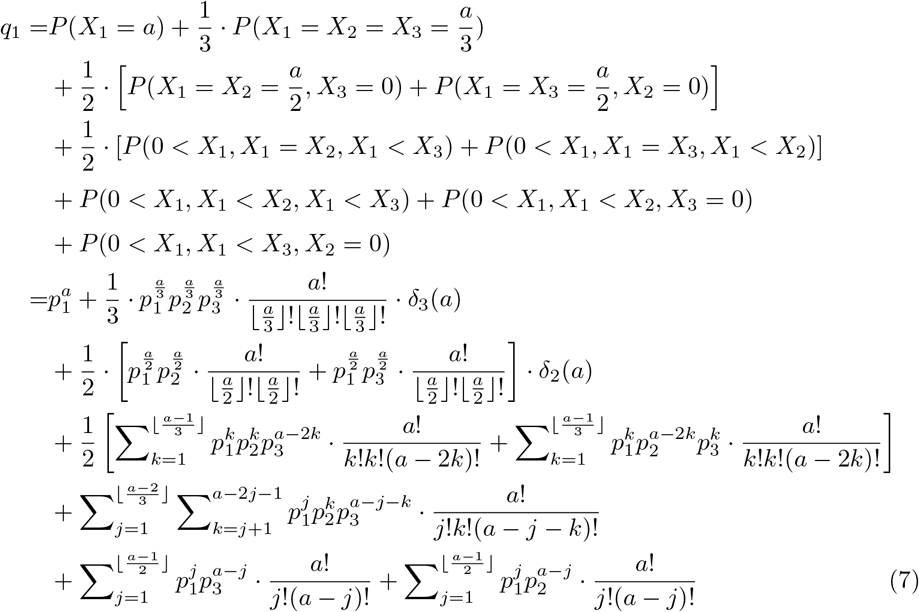

These examples show that it is possible to determine the probability of a father to fertilize a flower dependent on the locations of the trees as these define the probabilities to fertilize an ovule. Furthermore, it shows that the formula becomes more complicated when increasing the number of fathers and/or the number of ovules.

## B Appendix B

Figs. 11, 12, and 13 show the locations of the trees in the three different field studies used for validating our model.

**Fig 11.**
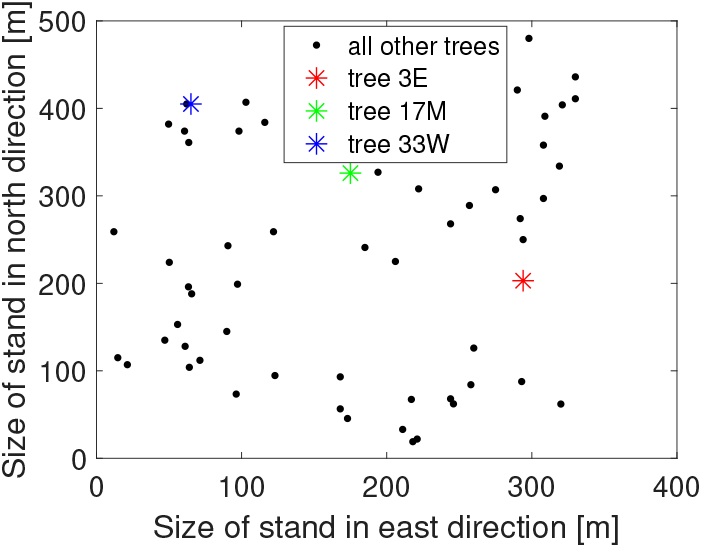
Locations of a stand of bur oaks in northern Illinois. Adapted from [2].

**Fig 12.**
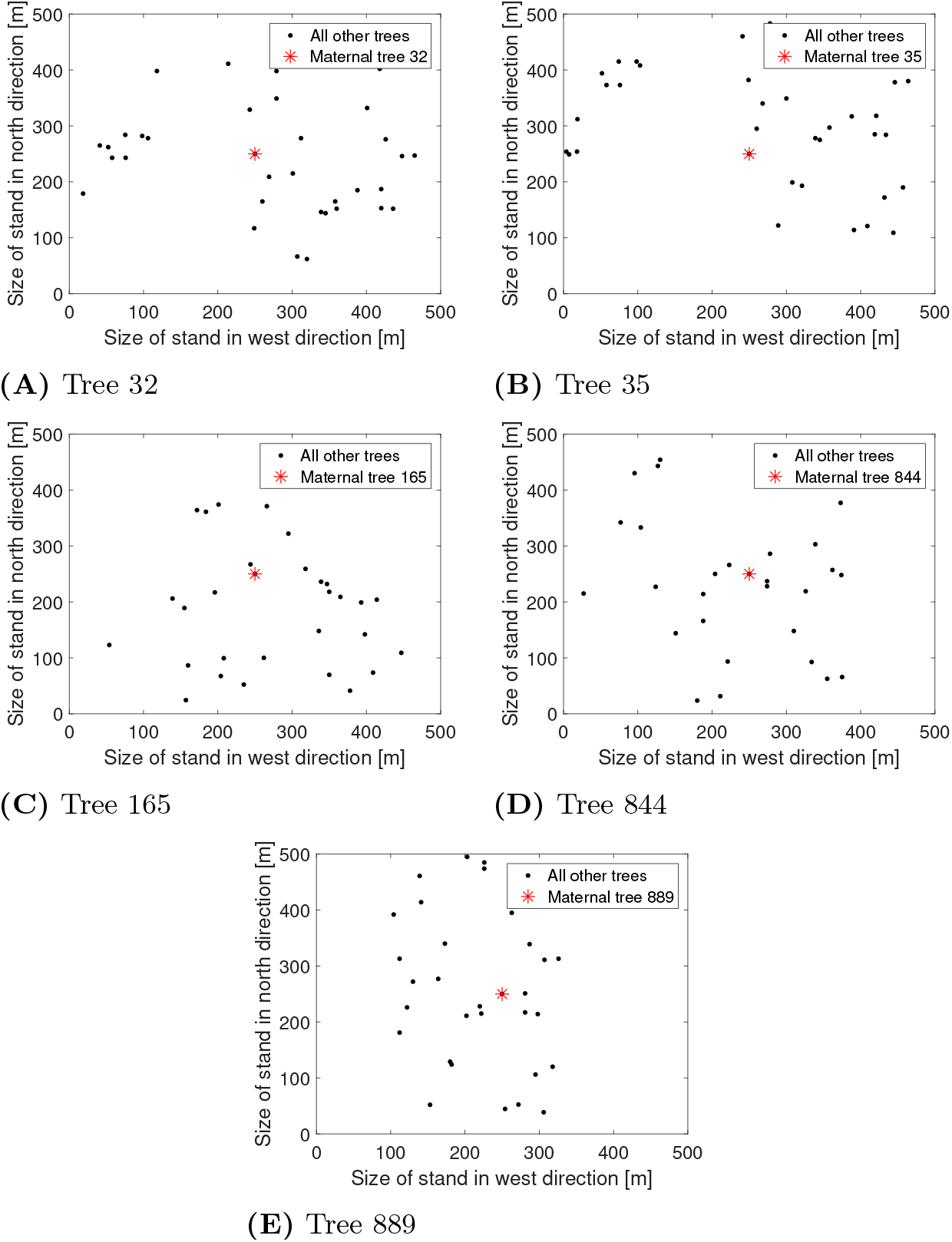
Locations for maternal trees 32, 35, 165, 844 and 889 and the surrounding trees. The pictures are adapted from [35].

**Fig 13.**
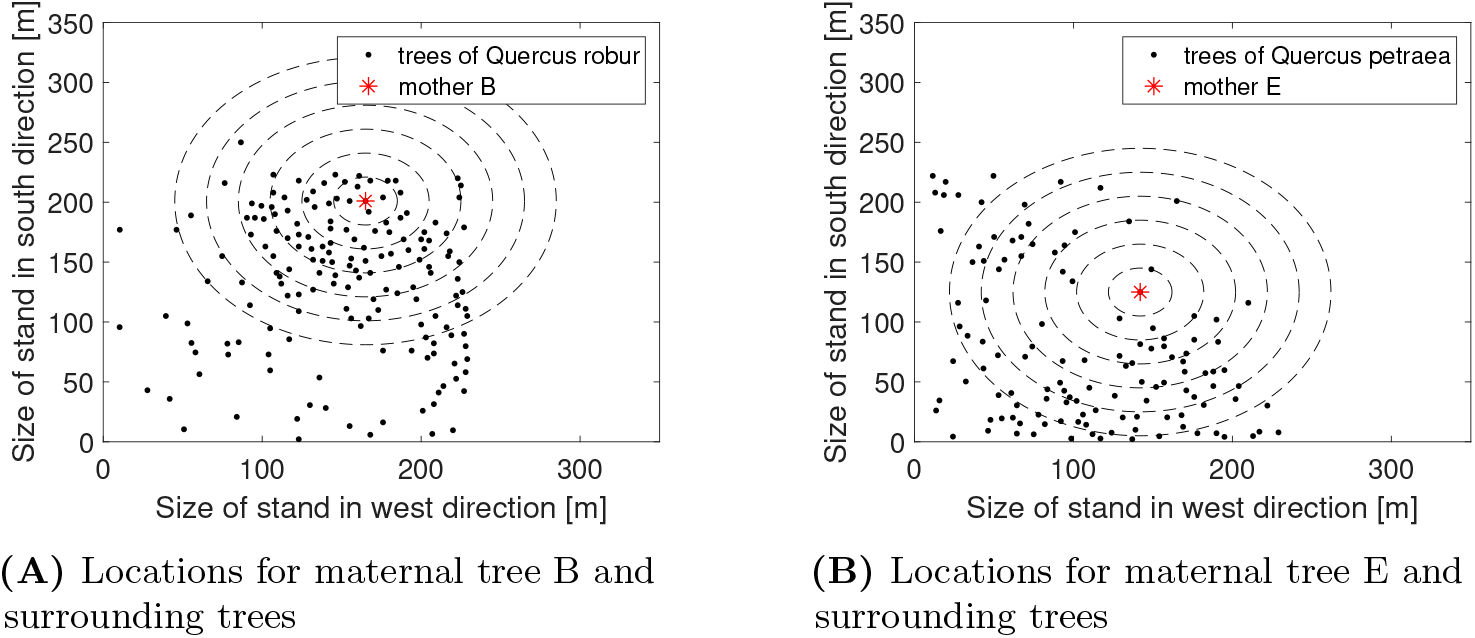
Figs 13A and 13B show the locations of oak trees. The rings surrounding the maternal trees indicate the distance classes. The locations are taken from [34].

